# Releasing potential power of terpene synthases by a robust precursor supply platform

**DOI:** 10.1101/105247

**Authors:** Guangkai Bian, Yichao Han, Anwei Hou, Yujie Yuan, Xinhua Liu, Zixin Deng, Tiangang Liu

## Abstract

Approximately 76,000 discovered makes terpenoids the largest family of natural products in nature with widespread applications. The wide-spectrum of structural diversity of the terpenoids were largely due to the variable skeletons generated by terpene synthases. The number of terpene skeletons found in nature, however, were so much more than those conceivably generated from known terpene synthases and the limited characterized terpene synthases also make no chance for some useful terpenoids overproduction in microbe. Here, we first demonstrated that the promiscuous synthases *in vivo* can produce more variable terpenoid products by converting precursors of different lengths (C_10_, C_15_, C_20_, C_25_). This discovery was prompted by the development of an efficient *in vivo* platform by combining the two promiscuous terpene synthases and three prenyltransferases to generate 50 terpenoids, at least 3 ring systems of which were completely new. Furthermore, protein engineering was further integrated to enhance product diversity. Clearly, the work is expected to dramatically reshape the terpenoid research by widening the flexibility of the terpene synthases for the fresh discovery or creation of the new terpenoid compounds by skeleton reframing.

## 1. Introduction

Nature is an excellent synthetic chemist, utilising two simple processes, i.e., mutations and combinations of biosynthetic genes, to create diverse natural products (Cane et al., 1998). By combining sets of genes, domains, and modules identified from different biosynthetic pathways or created by scientists in various ways, the concept of combinatorial biosynthesis has emerged to accelerate the discovery and expand the diversity of products (Kim et al., 2015; Tsoi and Khosla, 1995). This concept has been applied extensively to two large families of complex natural products, namely, polyketides and nonribosomal peptides (Khosla et al., 2014; Sieber and Marahiel, 2005; Weissman and Leadlay, 2005). Compared with these two families, very few reports have described the use of this strategy for terpenoids, the largest and most structurally diverse family of natural products. Recently, combinatorial biosynthesis has been proposed to generate diverse labdane-based diterpenoids (Andersen-Ranberg et al., 2016; Jia et al., 2016; Mafu et al., 2016). However, many other skeletons in the big terpenoid family remain to be explored.

The diversity of terpenoids arises from three steps (Fig. S1): (i) prenyltransferases (PTs) assemble the C_5_ precursors isopentenyl diphosphate (IPP) and dimethylallyl diphosphate (DMAPP) into polyisoprenoid diphosphates with various lengths, such as C10 geranyl diphosphate (GPP), C_15_ farnesyl diphosphate (FPP), C_20_ geranylgeranyl diphosphate (GGPP), and C_25_ geranylfarnesyl diphosphate (GFPP) (Kellogg and Poulter, 1997); (ii) terpene synthases cyclise these polyisoprenoid diphosphates to generate terpene products with a single ring or intricate multiple rings (Christianson, 2006); (iii) tailoring enzymes, including oxygenases, methyltransferases, acetyltransferases, and glycosyltransferases, add functional groups at different positions, further enhancing structural diversity. The possibility of obtaining linear products from the first step is very limited for the fixed length of C_5_ precursors, and the flexibility of the final products from the third step strongly depends on the skeletons from the second step. Thus, variability in the skeleton is mainly introduced through the cyclisation step. Biosynthesis by terpene synthases in this step has inherent advantages over chemical synthesis (Maimone and Baran, 2007); however, the biosynthetic repertoire identified to date only covers a small proportion of the terpene skeletons discovered, which only covers a small proportion of potential terpene skeletons through theoretical calculations (Tian et al., 2016). Additionally, most traditional methods to investigate terpene synthases only focus on their major functions under native conditions, ignoring their potential for different applications.

Combinatorial biosynthesis of terpenoid skeletons will be an effective solution when two prerequisites are satisfied: identification of terpene synthases that tolerate different lengths of isoprene-diphosphate as substrates, and generation of suitable polyisoprenoid diphosphate substrates for these enzymes. In this study, we successfully identified promiscuous terpene synthases in nature and developed a terpene-overproducing chassis to satisfy these two criteria. Six *Escherichia coli* variants, obtained by combining two terpene synthases and three PTs, generated 50 terpenoids. We also demonstrated that terpene synthases with substrate promiscuity are easily available in nature, and protein engineering of theses enzymes can be further incorporated in our strategy.

## 2. Materials and Methods

### 2.1 Materials

Linalool (51782, > 99%; Sigma-Aldrich, St. Louis, MO, USA), α-terpineol (30627, ≥ 90%; Sigma-Aldrich), pyrophosphate reagent (P7275; Sigma-Aldrich), and α-farnesene (F102425; J&K Scientific, Beijing, China) were used in this study. Kinetic assays were recorded on a Multimode Plate Reader Enspire (PerkinElmer, MA, USA). Proteins were purified using a Biologic DuoFlow Chromatography System (Bio-Rad, Hercules, CA, USA). Terpenoids were separated using a Dionex Ultimate 3000 UHPLC system (Thermo Scientific, Waltham, MA, USA). Optical rotations were measured on a PerkinElmer Model 341 polarimeter (PerkinElmer, USA). Single-crystal data were measured on an Bruker Kappa APEX-Duo CCD diffractometer (Bruker, MA, USA). ^1^H NMR and ^13^C NMR were performed on an Agilent 400 MHz or 600 MHz instrument (DirectDrive2; CA, USA).

### 2.2 Identification of class I terpene synthases

Class I terpene synthases were identified by screening all predicted proteins against a set of hidden Markov models constructed for each terpene synthase family. The model trained from the non-plant terpene cyclase alignment profile (cd00687 from the Conserved Domain Database) showed the best prediction coverage in our preliminary test data. The final identified proteins were manually curated on the basis of protein alignment and transcript assembly.

### 2.3 Phylogenetic analysis

Multiple sequence alignment was performed with MAFFT software (Katoh et al., 2002) using the local pair iterative refinement method. The following evolutionary analyses were conducted in MEGA7 (Kumar et al., 2016). The phylogenetic tree was inferred using the maximum likelihood method based on the Jones-Taylor-Thornton model. All positions containing gaps or missing data were eliminated. The final dataset included 211 positions.

### 2.4 Motif analysis

The active-site cavity was mapped according to existing crystal structures of fungal terpene synthases (PaFS, AtARS, and FsTDS) and refined alignment of all characterised fungal terpene synthases. The binding pocket of terpene synthases consisted of six α-helixes, i.e., C, D, F, G, H, and J. Strictly conserved and catalysis-related sites were excluded. Sequence logos of nonconserved residues in the active-site cavity were generated using the WebLogo server (http://weblogo.berkeley.edu/).

### 2.5 Molecular modeling

The three-dimensional model of the TC domain of FgMS was built using the SWISS-MODEL web server (https://www.swissmodel.expasy.org/). The crystal structure of the TC domain of PaFS from *Phomopsis amygdali* was selected as the template. The resulting homology structures were visualised using UCSF Chimera.

### 2.6 *In vitro* assays and kinetic measurements

To test the *in vitro* activities of FgMS, its mutants, and FgGS, reactions were conducted using 10 μM purified proteins, 100 μM substrates (GPP, FPP, GGPP, or GFPP), and 2 mM Mg^2+^ in 200 μL of 50 mM PB buffer (pH 7.6) with 10% glycerol at 30°C overnight. The sample was extracted with hexane (200 μL) and then analysed by GC-MS. For steady-state kinetics, 100-μL scale reactions were carried out in 50 mM Tris-HCl buffer (pH 7.6) with 10% glycerol and 50 μL pyrophosphate reagent. The concentration of the enzyme (FgMS or FgGS) were kept constant at 1 mg/mL with 2 mM Mg^2+^ as the only metal ion, while substrates (GPP, FPP, GGPP, and GFPP) were added at difference concentration ranging from 1 to 200 mM. Product assays were carried out by measuring the release of pyrophosphate (PPi), which recorded on an Enspire Multimode Plate Reader, using a previously described method (Agger et al., 2008).

### 2.7 Fermentation and purification of terpenoids

*E. coli* T7 (harbouring pMH1, pFZ81, and pGB310), T8 (harbouring pMH1, pFZ81, and pGB311), T9 (harbouring pMH1, pFZ81, and pGB312), T10 (harbouring pMH1, pFZ81, and pGB313), and T11 (harbouring pMH1, pFZ81, and pGB314) were cultivated in 2-L flasks containing 1 L LB medium at 37°C with 100 mg/L ampicillin (AMP), 50 mg/L KAN, and 34 mg/L chloramphenicol (CM). When the OD**_600_** reached 0.6–0.8, 0.1 mM IPTG was added to the cultures, protein expression was induced at 16°C for 18 h, and the strains were then cultivated at 28°C for an additional 3 days. The cell cultures were extracted twice with an equal volume of hexane. The organic layer was concentrated using a rotary evaporator, and terpenoids were purified using a Dionex Ultimate 3000 UHPLC system.

### 2.8 Detection of terpenoids

Terpenoids were extracted by hexane and detected by GC-MS (Thermo TRACE GC ULTRA combined with a TSQ QUANTUM XLS MS). The samples were injected into a TRACE TR-5MS (30 m × 0.25 mm × 0.25 um). The oven temperature was set at 80°C for 1 min, increased to 220°C at a rate of 10°C/min, and held at 220°C for 15 min. The injector and transfer lines were maintained at 230°C and 240°C, respectively. Product structures were determined by comparison with authentic standards, comparison with mass spectra data from the NIST library, and NMR analysis.

## 3. Results

### 3.1 Terpene synthases with substrate promiscuity

To predict promiscuous terpene synthases from sequences, we collected sequences of terpene synthases in fungi, a major source of terpenoids that has not yet been fully explored, and constructed a phylogenic tree. Fifty-one genetically or biochemically characterised terpene synthases (Additional Data Table S1), including mono-, sesqui-, di-, and sesterterpene synthases, were mainly divided into five clades (Fig. 1a). One of them, clade III, was the most interesting. Notably, as of the writing of this manuscript (August 2016), all characterised di- and sesterterpene synthases in fungi were enriched in clade III rather than widely distributed, while diterpene synthases were dispersed in the phylogenic trees of bacterial terpene synthases (Dickschat, 2016; Yamada et al., 2015). Thus, enzymes in this clade are more likely to generate large complex terpene skeletons (Fig. 1b). More importantly, in this clade, three enzymes are able to catalyse the cyclisation of multiple polyisoprenoid diphosphate substrates *in vitro* (Chen et al., 2016; Chiba et al., 2013; Qin et al., 2016). Although the substrate specificities of other enzymes have not been investigated to date, we speculated that the broad substrate range could be a common property of this type of terpene synthase. For these two reasons, we selected enzymes in clade III as candidates for combinatorial biosynthesis.

**Fig. 1.**
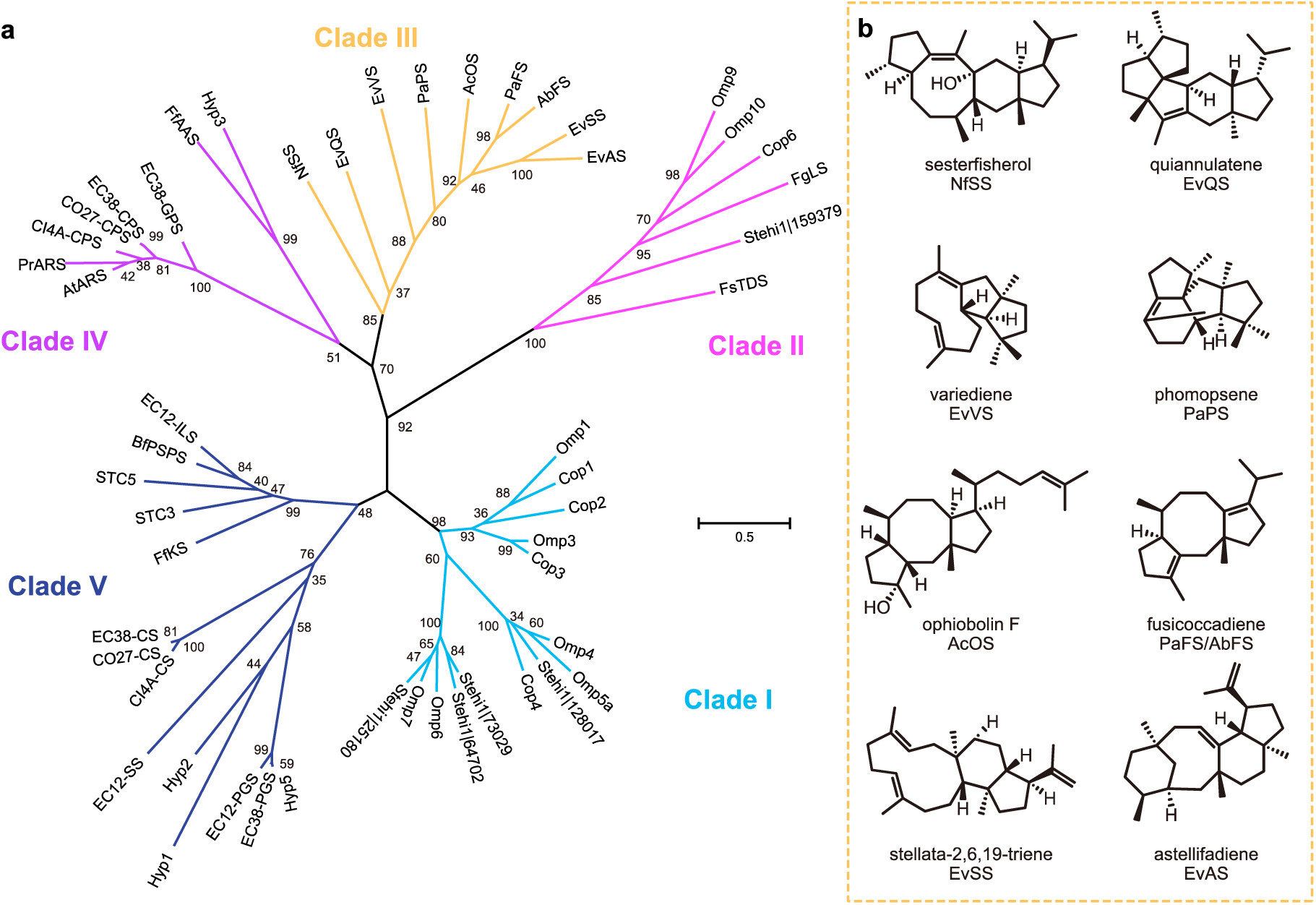
Clades of characterised fungal class I terpene synthases. (a) Phylogenetic tree of 51 experimentally characterised fungal terpene synthases. The enzyme abbreviations are described in the Additional Data Table S1. Branch length indicates the substitutions per site. Branches are labeled with the percentage of 1,000 bootstrap replicates. (b) Products of terpene synthases in clade III in previous studies.

From the collection of genomes we sequenced in our lab, we found two clade III terpene synthases in an endophytic fungus *Fusarium graminearum* J1-012, a well-known pathogen causing Fusarium head blight. These two enzymes (i.e., FgJ07578 and FgJ07623 in our annotated genome) were selected as candidates because no similar clade III enzymes of them have been characterised before. FgJ07578 is a chimeric protein consisting of an N-terminal terpene cyclase (TC) domain and a C-terminal PT domain; in contrast, FgJ07623 only contains a TC domain of 316 amino acids. Notably, no PT exists near FgJ07623, and the biosynthetic genes in its cluster are not typically related to terpenoid synthesis (Fig. S2).

### 3.2 Validating broad substrate specificities

These two enzymes (FgJ07578 and FgJ07623) were overexpressed in *E. coli,* purified, and characterised biochemically *in vitro.* Surprisingly, in the absence of IPP, both FgJ07578 and FgJ07623 could directly catalyse the conversion GPP, FPP, GGPP, and GFPP to generate multiple products *in vitro*, as detected by gas chromatography mass spectrometry (GC-MS), exhibiting unprecedentedly broad substrate specificities for polyisoprenoid diphosphates (Fig. 2).

**Figure 2.**
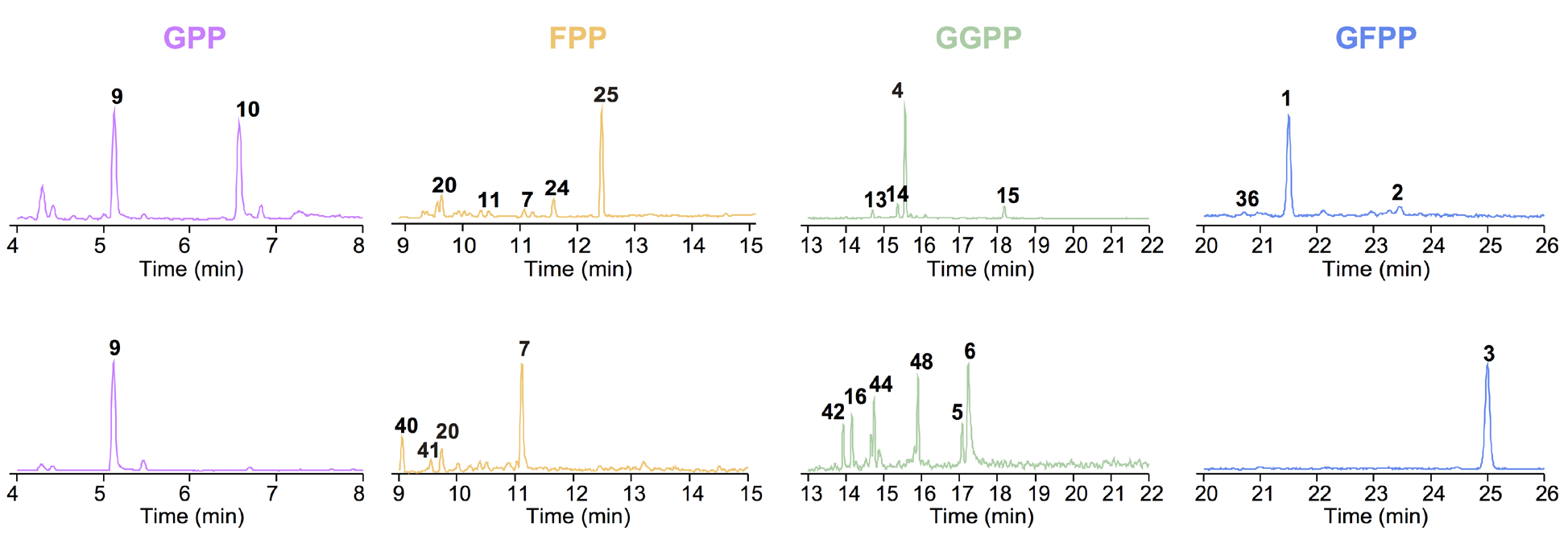
*In vitro* characterisation of FgJ07578 and FgJ07623. MS analysis of terpenoid products from *in vitro* assays of FgJ07578 (top) and FgJ07623 (bottom) with GPP, FPP, GGPP, and GFPP, respectively, in the absence of IPP.

Because isoprenoid diphosphate substrates could be elongated by the PT domain of FgJ07578 itself, we also conducted *in vitro* experiments with the addition of IPP for FgJ07578. When DMAPP and IPP were used as substrates, mono-, sesqui-, di-, and sesterterpenes were all detected through *in vitro* reactions, indicating that the PT domain of FgJ07578 could generate multiple polyisoprenoid diphosphates from C_10_ to C_25_ *in vitro* (Fig. S3). In addition, when GPP, FPP, or GGPP was incubated with the addition of IPP as substrates (Fig. S3), both terpenes with corresponding lengths and terpenes with longer lengths were generated, suggesting that polyisoprenoid diphosphate substrates could be not only directly cyclised by the TC domain but also first elongated by the PT domain to form longer substrates.

We then performed kinetic analyses to understand the turnover rates and specificities quantitatively. When substrates reaching their saturating concentrations, FgJ07578 exhibited higher turnover rates for GGPP and GFPP (almost equally), and FgJ07623 exhibited the highest turnover rate for GGPP. However, both FgJ07578 and FgJ07623 will prefer GPP as the better substrate if different substrates were given, because of their highest specificity constants (*k_cat_*/*K_m_*) for GPP. Notably, the specificity constants towards FPP, GGPP, and GFPP were similar for these two enzymes, with their differences being less than 1.5-fold change (Table 2). This similarity provided a basis for engineering to exploit substrate promiscuity by redistributing substrates *in vivo*.

**Table 1.**
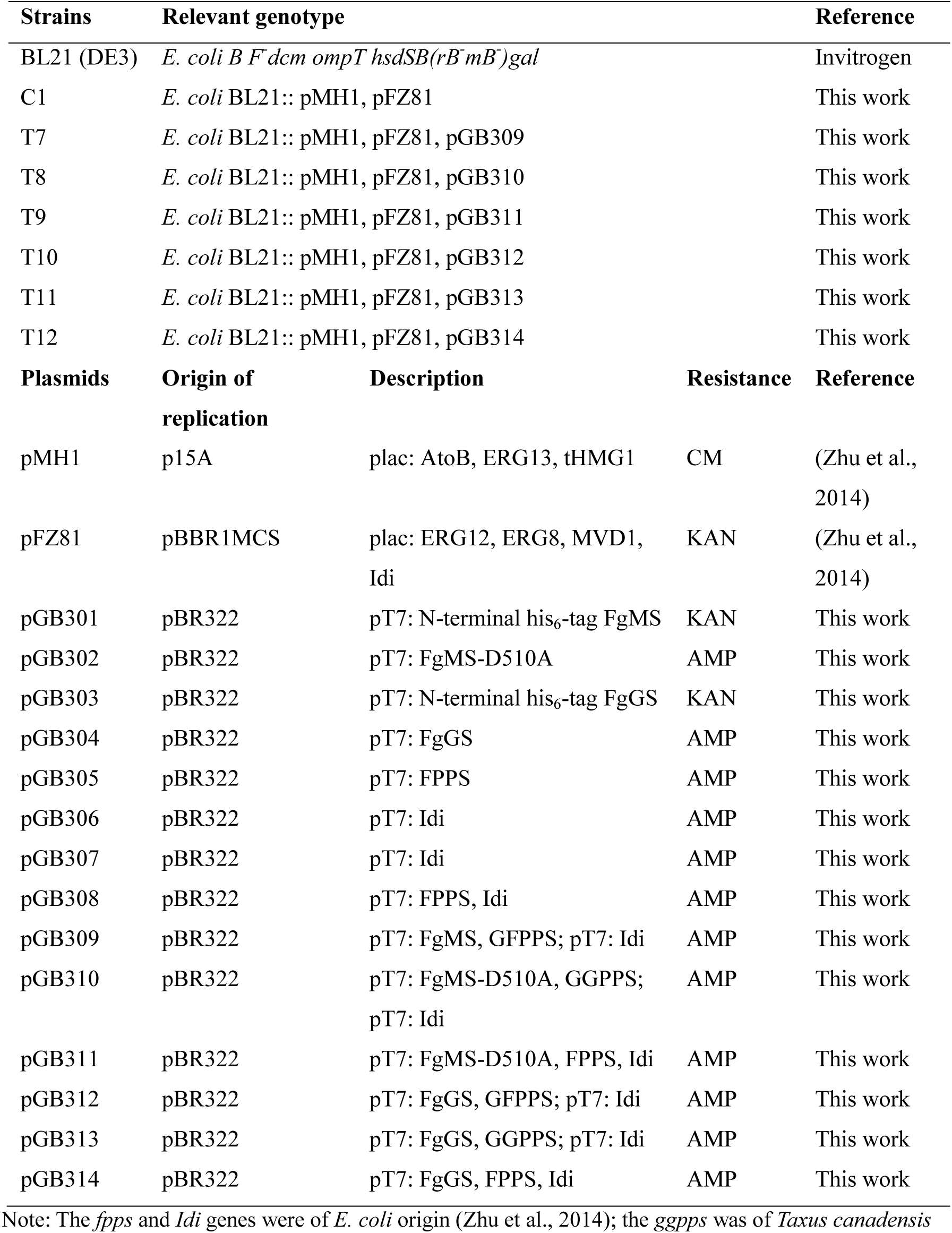
Strains and plasmids used in this research.

**Table 2.**
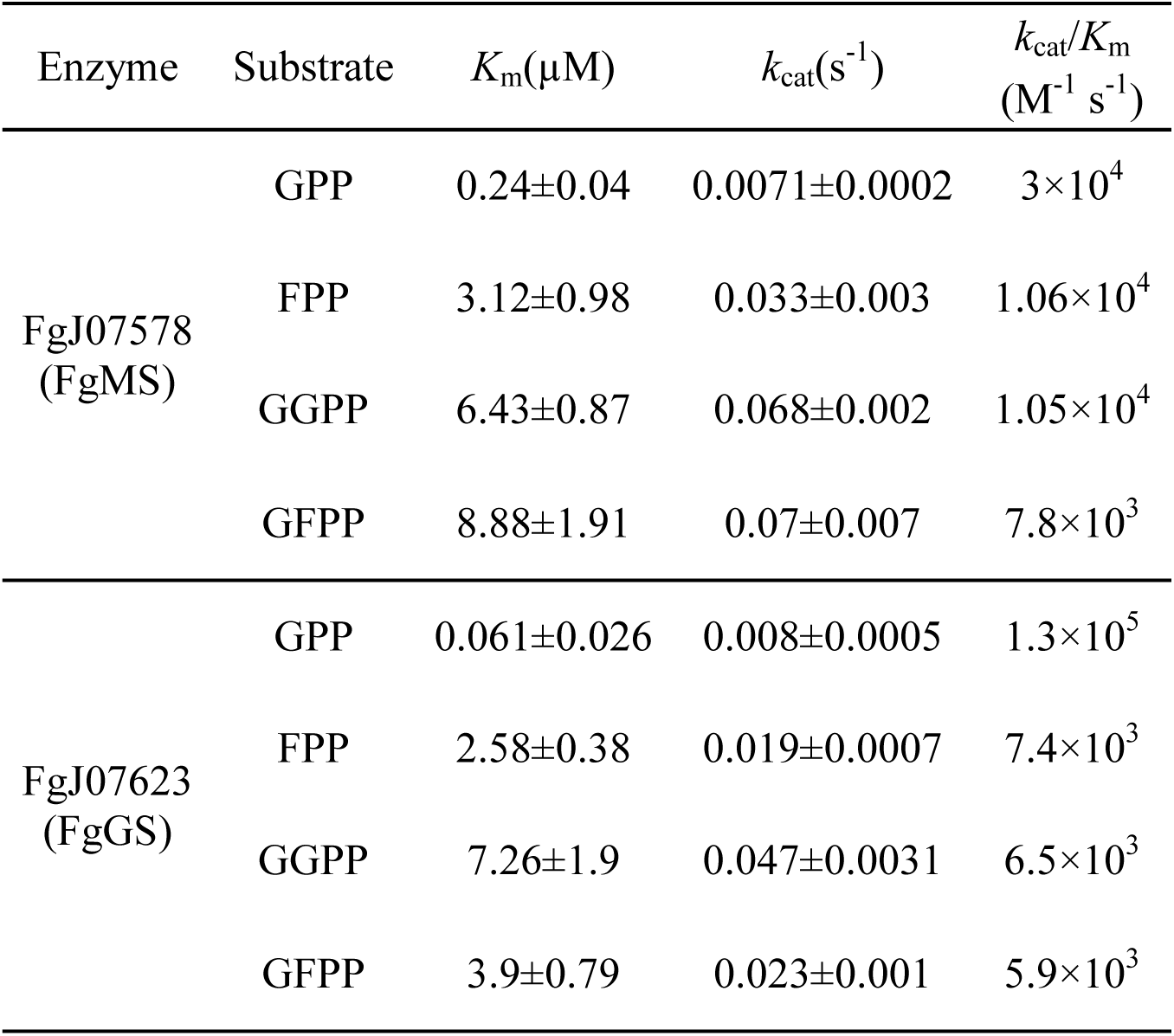
Catalytic activity of FgMS and FgGS.

### 3.3 Combinatorial biosynthesis of terpenoid skeletons

To realise the full potential of these promiscuous terpene synthases, we developed a platform consisting of three parts: a terpene-overproducing chassis, a PT library, and the promiscuous terpene synthases of interest. First, we built an *E. coli* strain as a terpene-overproducing chassis based on previous design principles (Zhu et al., 2014). Specifically, on the basis of data from *in vitro* reconstitution, proteomics, and metabolomics, the entire mevalonate pathway (AtoB, ERG13, tHMG1, ERG13, ERG8, MVD1, and Idi) was appropriately constructed to provide sufficient C5 precursors (Fig. 3a). This strategy has successfully been applied to overproduction of farnesene (Zhu et al., 2014), lycopene (Zhu et al., 2015), astaxanthin (Ma et al., 2016), and taxadiene (unpublished data). Second, efficient PTs, including FPPS, GGPPS, and GFPPS from previous reports (Ajikumar et al., 2010; Furubayashi et al., 2015; Zhu et al., 2014), were selected and codon-optimised to construct a library. GPPS was not included because of the relatively limited possible conformations of monoterpenes. Third, we combined the two terpene synthases with these three PTs to form six variants from T7 to T12 (Fig. 3b). In *E. coli* T8 and T9, the PT domain of FgJ07578 was inactivated by replacing a key catalytic residue, aspartic acid, with alanine (D510A) to block substrate flux towards GFPP.

**Fig. 3.**
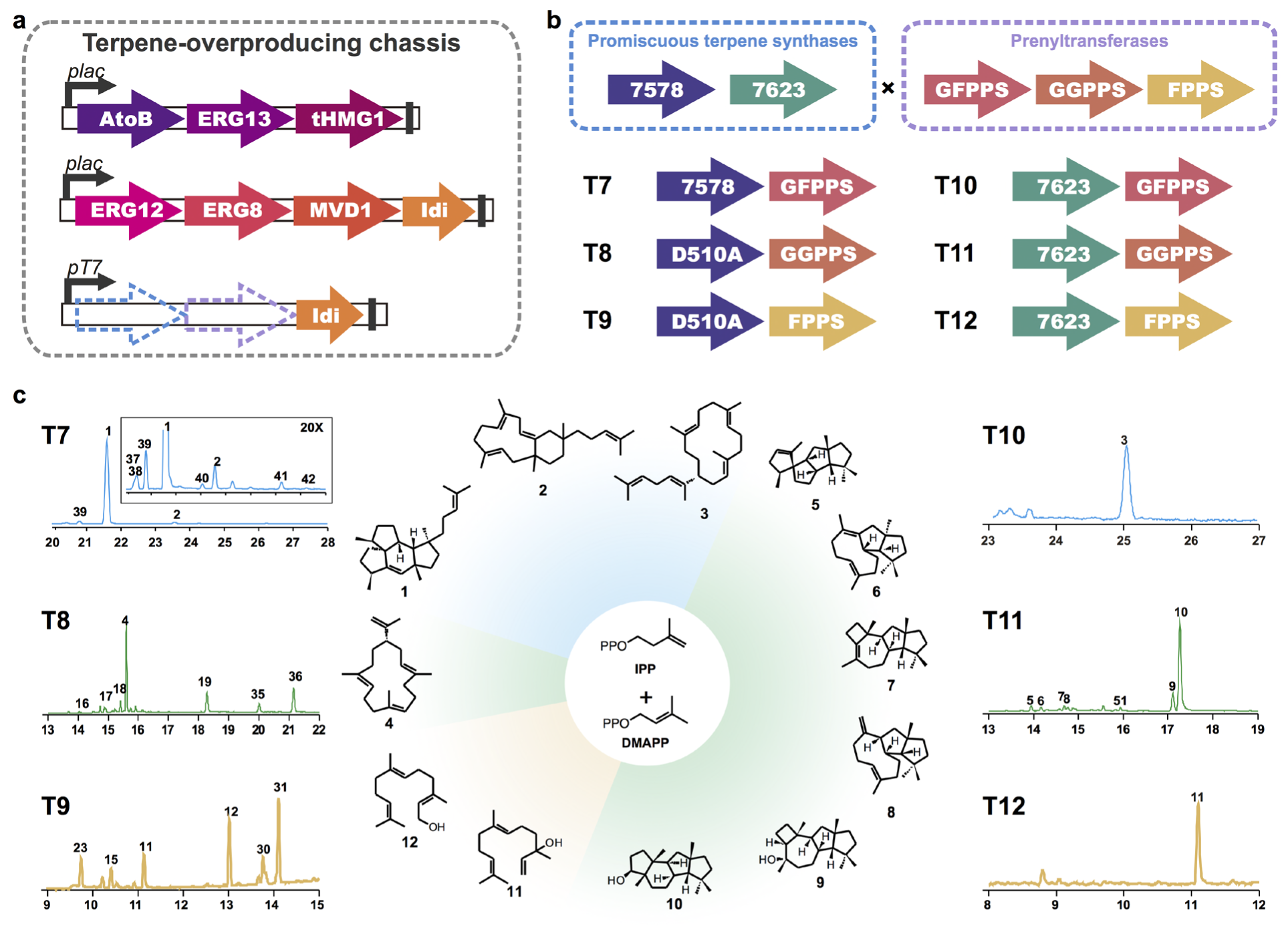
Accessing diverse complex terpenoid skeletons using combinatorial biosynthesis. **(a)** A terpene-overproducing chassis in which the entire mevalonate pathway was introduced. (**b**) Combinations of two terpene synthases and three prenyltransferases. 7578: FgJ07578; 7623: FgJ07623; D510A: the D510A mutant of FgJ07578. (c) Fermentation results of six *E. coli* strains. All the structures shown were identified by NMR analysis.

GC-MS analysis revealed that 50 different terpenoids were produced by these six strains, including nine sesterterpenoids, 24 diterpenoids, and 17 sesquiterpenoids (Fig. 3c, Figs. S4a and S5). Twelve products (**1** to **12**) were separated followed by nuclear magnetic resonance (NMR) analyses to determine their chemical structures.

Three sesterterpenoids were subjected to NMR analyses, two of which were new precursors of natural products with pharmaceutical activities. Specifically, two sesterterpenoid products (**1** and **2**) in *E. coli* T7 were identified as a 5-5-6-5 tetracyclic compound and an 11-6 bicyclic compound, respectively. Compound 1 is the sesterterpene precursor of mangicols, of which mangicol A and mangicol B exhibit anti-inflammatory activity (Renner et al., 2000). Compound **2** is the sesterterpene precursor of variecolin, which has been shown to inhibit angiotensin II receptor (Hensens et al., 1991) and act as an immunosuppressant (Fujimoto et al., 2000). The identification of these skeletons could fill gaps between biosynthetic repertoire and known natural products. The cyclisation mechanisms of these two compounds were summarised in Fig. S6a according to previous classification (Ye et al., 2015). Besides these two compounds, the GC-MS data (Fig. 3c and Fig. S5) of another six sesterterpenoids (**37** to **42**) detected from *E. coli* T7 cannot be matched with existing sesterterpenoids data in the literatures, suggesting that new structures could be discovered. The only sesterterpenoid product (**3**) in *E. coli* T10 was determined to be *2E*-alpha-cericerene; this compound is also a sesterterpene product of the EvVS variant (Qin et al., 2016), which shares only 40% sequence identity with FgJ07623.

From structural elucidation of diterpene products, three entirely new ring systems were identified. The diterpenoid main product (**4**) in *E. coli* T8 was found to be cembrene A, a simple skeleton with a large 14-membered ring that has been reported to be a byproduct of the bacterial diterpene synthase DtcycB (Meguro et al., 2013). In addition, some of its side diterpene products, such as **18** and **19**, were predicted to have the same skeleton as cembrene A on the basis of mass spectra data from the NIST library (Figs. S4a and S5). To the best of our knowledge, this was the first fungal gene found to generate this type of C20 single-large-ring skeleton. The skeletons of diterpenoids **5** to **10** in *E. coli* T11 all possessed two five-membered rings, which were formed in the first two ring-closure steps (Fig. S6b). Compound 5 was identified as a 5-5-5-5 membered tetracyclic diterpene. Compounds **6** and **8** were identified as 5-5-9 membered tricyclic diterpenes, which differed in the positions of double bonds. The relative configuration of compound **6** was matched with variediene (Qin et al., 2016). Compound **7** was identified as a 5-5-7-4 tetracyclic diterpene. Compounds **9** and **10** were identified as 5-5-7-4 and 5-5-6-5 tetracyclic diterpene alcohols without any double bonds, respectively. To the best of our knowledge, the 5-5-5-5, 5-56-5, and 5-5-7-4 tetracyclic skeletons are unprecedented. The absolute configuration of **10** was then determined by single-crystal X-ray diffraction analysis (Additional File).

Sesquiterpenoids identified were often linear products or alcohols. Two sesquiterpenoid main products (**11** and **12**) in *E. coli* T9 were validated as trans-nerolidol and *2E, 6E*-farnesol, two linear alcohols. *E. coli* T12 also produced *trans*-nerolidol as the main product, which is one of the few overlapping products of these two terpene synthases (Fig. S7).

According to existing natural products mangicol and variecolin, the new products 1 and 2 were designated mangicdiene and variecoltetraene respectively. Because no reference could be given to new products **5** and **7** to **10**, they were designated GJ1012 A to E, respectively; thus, these two terpene synthases (FgJ07578 and FgJ07623) were designated *F. graminearum* mangicdiene synthase (FgMS) and *F. graminearum* GJ1012 synthase (FgGS), respectively.

### 3.4 Other promiscuous terpene synthases in this clade

To test whether the substrate promiscuity is a widespread feature of this kind of terpene synthases, we performed large scale genome mining. In UniProt database, 2088 proteins from 208 fungal genomes were predicted to be class I terpene synthases. According to our phylogenic analysis, 321 (15.4%) of them could be further classified as clade III terpene synthases in 109 different species (Additional Data Table S2). The result indicates clade III terpene synthases are easily available for application of our strategy.

### 3.5 Incorporating protein mutagenesis

To further enhance product diversity, we performed site-direct mutations of the promiscuous FgMS as an example to integrate protein engineering into our framework. We first investigated how the diverse sequence features of natural terpene synthases led to functional differentiation in evolution. At least 13 noncatalytic sites within the pocket were diverse in fungi (Table S1). For convenience, we named these specificity-determining sites (SDSs) 1–13, corresponding to residue numbers 65, 69, 85, 88, 89, 159, 168, 191, 219, 222, 307, 311, and 314 in FgMS. In the terpene cyclisation reaction, the aromatic residues phenylalanine (F), tryptophan (W), and tyrosine (Y) not only stabilise carbocation centres through cation-π interactions but also provide steric hindrance, restricting the active-site contour (Li et al., 2014); therefore, we highlighted the SDSs where aromatic residues occurred in the sequence logo for each clade of the phylogenetic tree (Fig. 4a). Aromatic residues in SDSs 5, 6, and 13 were relatively conserved in all fungal terpene synthases; however, other SDSs with abundant aromatic residues seemed specific to distinct clades, e.g., they were enriched in SDSs 1, 8, and 9 in clade III and in SDSs 2, 11, or 12 in other clades. Thus, based on our observations of natural evolution, our protein engineering strategy focused on the interchange between aromatic and nonaromatic residues for each SDS in FgMS (Fig. 4b).

**Figure 4.**
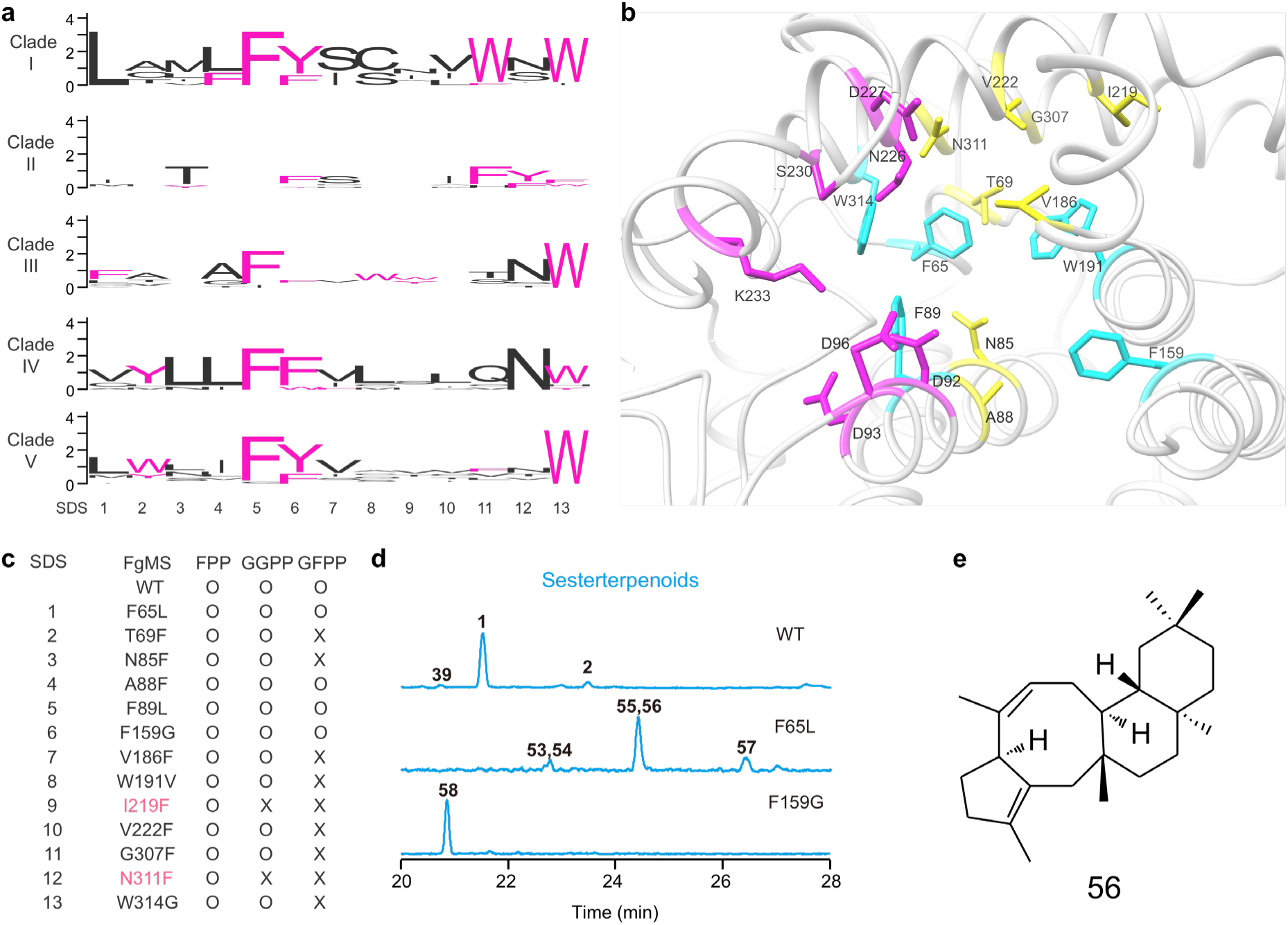
Protein engineering by interchange between aromatic and nonaromatic residues. (a) Sequence logos of specificity-determining sites (SDSs) for each clade in the phylogenetic tree. SDSs 1-13 in these logos correspond to residue numbers 65, 69, 85, 88, 89, 159, 168, 191, 219, 222, 307, 311, and 314 in FgMS. Aromatic residues are highlighted. (b) Three-dimensional model of the active-site cavity of FgMS. Metal-binding catalytic motifs are indicated in magenta, aromatic SDSs are indicated in cyan, and nonaromatic SDSs are indicated in yellow. (c) Substrate ranges of FgMS mutants. O, maintain catalytic activity; X, loss catalytic activity. (d) New products can be generated by FgMS mutants. WT, wild-type FgMS. (e) The chemical structure of compound 56.

Our preliminary cell-free incubation experiments showed that when nonaromatic residues of SDSs in FgMS were replaced by F, W, or Y, the mutants generated the same products, but with different product ratios, i.e., substitutions of different aromatic residues did not alter the product compositions of FgMS. Therefore, we only used the mutants replaced by F to represent the interchange from a nonaromatic residue to an aromatic residue. All mutants of interchange between aromatic and nonaromatic residues for SDSs maintained their catalytic activity, at least for FPP (Additional Data Table S3 and Fig. 4c). Nine of the mutants exhibited narrower substrate ranges, and two of them even lost catalytic activity for GGPP (Fig. 4c). These phenomena also implied that interchange between aromatic and nonaromatic residues could be utilised to broaden the substrate ranges of other terpene synthases with narrow ones. From a homolog model view, when the nonaromatic hydrophobic residues were replaced by aromatic hydrophobic residues in the middle or bottom of the binding pocket, the overall pocket volume will be greatly reduced disabling the catalysis of longer substrates. Position A88 was an exception because it did not point to the center of the binding pocket (Fig. 4b). Comparing the sequences of characterised diterpene synthases and sesterterpene synthases could also offer similar explanations. PaFS cannot accept GFPP and it possesses more aromatic residues in their binding pockets (at SDS 1, 5, 6, 8, 9, and 13) (Table S1).

For the majority, the product profiles were redistributed, thus we could exploit different mutants to enhance production of certain products (Additional Data Table S3). More importantly, new products could be generated in this way to further enhance the terpene skeleton diversity. For examples, six new sesterterpenoids (**53 to 58**) emerged from mutants F65L and F159G (Fig. 4d and Fig. S5). Compound **56** was isolated and identified as a new 5-8-6-6 tetracyclic sesterterpene (Fig. 4e), which has the same skeleton as asperterpenol(Xiao et al., 2013), a fungal metabolite showing inhibition on acetylcholinesterase (acetylcholinesterase inhibition is one of current strategies to Alzheimer’s disease). In summary, incorporating protein engineering methodology boosted the performance of the combinatorial biosynthesis strategy.

## 4. Discussion

Our findings suggested that a relative small set of building blocks was sufficient to generate skeletally diverse terpenoids, demonstrating the elegance of terpene biosynthesis in nature. Although we did not test the physiological roles of the promiscuous terpene synthases (because they were silenced in the native fungi under our laboratory conditions), the implement of their multiple biosynthetic functions in living cells suggests the promiscuity of these enzymes could serve as regulation of cell metabolism. Similar examples exist in nature. Plants have utilized promiscuous terpene synthases to regulate emission of different flavors: controlled by targeting signals, nearly identical terpene synthases that can convert both GPP and FPP are delivered to different subcellular locations where different substrates are available, to produce C_10_ linalool and C_15_ nerolidol (Aharoni et al., 2004; Nagegowda et al., 2008), respectively. It is plausible that organisms preserve their promiscuous activity through evolution to increase fitness for survival because the more products that are generated, the more likely the organism is to produce chemicals with useful biological activity (Fischbach and Clardy, 2007).

Our discoveries and combinatorial biosynthesis strategy accelerated the entire process for discovery and creation of terpenoids. We demonstrated that terpene synthases with substrate promiscuity were widespread. Hundreds of protein candidates with similar property existed in current database. In addition, engineering the residues in the binding pocket can be further incorporated to access more scaffolds. Notably, the substrate promiscuity was not unique in di/sesterterpene synthases, which were concentrated in clade III fungal terpene synthases. We focused on this kind of enzymes because they could be utilized to generate relatively complex compounds with one or more rings. The polycyclic skeletons extensive in di- and sesterterpenoids offer a variety of diverse structures, promoting numerous pharmaceutical applications (Stockdale and Williams, 2015), e.g. diterpenoids Taxol (paclitaxel) used in cancer therapy. Many skeletons, e.g. compound **1**, **2**, and **56**, we produced in this study are precursors of natural products with important pharmaceutical activities (Fig. S4b). The metabolic engineering platform helps us obtain a sufficient amount of these scaffolds, not only enabling the elucidation of their chemical structures, but also being a start for industrial application.

Our work greatly expanded the biosynthetic repertoire of terpene skeletons, i.e., the possibilities of upstream in the entire biosynthetic pathway. Combinatorial biosynthesis can also be applied to downstream modifications via promiscuous tailoring enzymes (Mafu et al., 2016). To further incorporating downstream tailoring enzymes, we recommended to transfer our strategy to engineered terpene-producing *Saccharomyces cerevisiae.* This is because heterologous expression of downstream decorating pathway in *E. coli* is still challenging, although some problems are progressively understood and solved (Biggs et al., 2016). Given that very limited terpenoids have been commercially generated through biosynthesis (Leavell et al., 2016), once multistep combinations are achieved, the fermentative production will cover much more desired terpenoids.

## 5. GenBank accession number

The DNA and protein sequences of FgJ07578, FgJ07623 and GFPPS described here are deposited in GenBank under accession numbers KY462789, KY462790 and KY462792.

## Acknowledgments

We thank Dr. Yuanzhen Liu (Wuhan University) and Dr. Zhongzheng Yang (Wuhan Institute of Biotechnology) for suggestions in NMR data analysis. This research was supported by the funding from J1 Biotech Co., Ltd. and grants from 3551 Optics Valley talent program for Prof. Zixin Deng, Science and Technology Project of Hubei Province (2013ACA011), the 973 Project of National Program on Key Basic Research (2012CB721000), and the Young Talents Program of National High-level Personnel of Special Support Program (The “Ten Thousand Talent Program”).

## Author contributions

G. B., Y. H., Z. D., and T. L. conceived of the project, designed the experiments, and wrote the manuscript; G. B., Y. Y., and X. L. performed *in vivo* and *in vitro* experiments; Y. H. performed bioinformatics analyses; A. H. performed NMR analyses; X. L. constructed and characterised the mutant library. Z. D. and T. L. managed the project.

## Competing financial interests

Two patent applications based on the results in this study have been filed by J1 Biotech Co., Ltd.

